# Experimental Evolution Reveals Redox State Modulates Mycobacterial Pathogenicity

**DOI:** 10.1101/2021.12.23.474061

**Authors:** Zheng Jiang, Zengfang Zhuang, Kaixia Mi

## Abstract

Understanding how *Mycobacterium tuberculosis* has evolved into a professional pathogen is helpful in studying its pathogenesis and for designing vaccines. We investigated how the evolutionary adaptation of *M. smegmatis* mc^2^51 to an important clinical stressor H_2_O_2_ allows bacteria undergo coordinated genetic mutations, resulting in increased pathogenicity. Whole-genome sequencing identified a mutation site in the *fur* gene, which caused increased expression of *katG*. Using a Wayne dormancy model, mc^2^51 showed a growth advantage over its parental strain mc^2^155 in recovering from dormancy under anaerobic conditions. Meanwhile, the high level of KatG in mc^2^51 was accompanied by a low level of ATP, which meant that mc^2^51 is at a low respiratory level. Additionally, the redox-related protein Rv1996 showed different phenotypes in different specific redox states in *M. smegmatis* mc^2^155, mc^2^51, *M. bovis* BCG and *M. tuberculosis* mc^2^7000. In conclusion, our study shows that the same gene presents different phenotypes under different physiological conditions. This may partly explain why *M. smegmatis* and *M. tuberculosis* have similar virulence factors and signaling transduction systems such as two-component systems and sigma factors, but due to the different redox states in the corresponding bacteria, *M. smegmatis* is a nonpathogen, while *M. tuberculosis* is a pathogen. As mc^2^51 overcomes its shortcomings of rapid removal, it can be potentially developed as a vaccine vector.

## Introduction

Tuberculosis (TB) is caused by the pathogen *Mycobacterium tuberculosis* and remains a public health threat, resulting in 1.4 million deaths in 2020 (WHO, 2021). The prevalence of multidrug-resistant *M. tuberculosis* and the rising cases of co-infection with HIV increase this health concern. The World Health Organization (WHO) has estimated that a quarter of the world’s population is infected with *M. tuberculosis* (WHO, 2021). This latent state may be extended as long as the life of the infected host, but, unfortunately, the reactive rate is approximately 5-10% of infected individuals (Flynn and Chan, 2001). TB is treated with chemotherapy, and the latent state of mycobacteria prolongs the time of treatment, which is one of the causes for the development of mycobacterial resistance.

As one of the world’s most successful human pathogens, *M. tuberculosis* has evolved elegant strategies to escape the immune defensive system of the host. For example, D’Arcy et al. observed that *M. tuberculosis*-containing phagosomes do not fuse with the lysosome inside infected macrophages (Armstrong and Hart, 1971; Brown et al., 1969). Several studies have indicated that *M. tuberculosis* as an intracellular pathogen is partially due to its ability to survive and persist in macrophages, in hostile environments with oxidative stress, in low pH, and under starvation, and other stresses (Cohen et al., 2018; Nauseef, 2019). When *M. tuberculosis* infects macrophages, mycobacteria must overcome exogenous reactive oxygen species (ROS), a classical innate defense mechanism against infection (Pieters, 2008). In addition, during latency, *M. tuberculosis* continues to be exposed to oxidative stress, and thus the accumulation of mutations caused by oxidative DNA damage is predicted as a potential risk for resistance to antibiotics (Ford et al., 2011). Clinical investigation has shown that cells from TB patients produce less ROS than those of healthy individuals (Jaswal et al., 1992; Kumar et al., 1995). In addition, ROS can also be associated with the treatment of TB (Hu et al., 2021; Piccaro et al., 2014). To avoid ROS attack evolved detoxified system is essential for *M. tuberculosis* survival, persistence and subsequent reactivation (Piccaro et al., 2014).

As a mycobacterial model of *M. tuberculosis, M. smegmatis* is contributed to understanding the functions of mycobacteria (Aldridge et al., 2012; Gray et al., 2016; Kieser et al., 2015). The essential genes in *M. tuberculosis* have a corresponding ortholog gene in *M. smegmatis* (Dragset et al., 2019; Judd et al., 2021). *M. smegmatis* and *M. tuberculosis* H37Rv share 2547 mutually orthologous genes (Dragset et al., 2019). At least under certain conditions, *M. tuberculosis* and *M. smegmatis* have similar growth mechanisms. The assumed virulence factors identified in *M. tuberculosis* such as PhoPR and DosR/S/T (Gonzalo-Asensio et al., 2014; Mehra et al., 2015) are also present in *M. smegmatis*. However, compared to virulent *M. tuberculosis*, *M. smegmatis* is a nonpathogenic mycobacterium. *M. tuberculosis* can persist in the infected host, while the host can quickly remove *M. smegmatis* (Anes et al., 2003; Anes et al., 2006). Considering that *M. tuberculosis* with high resistance to hydrogen peroxide (H_2_O_2_) persists in the lungs, while *M. smegmatis* with lower resistance to H_2_O_2_ exists in the soil, we hypothesized that the redox state of *M. tuberculosis* and *M. smegmatis* may adapt to the corresponding redox environment. The differences between the two bacteria are due to their corresponding degree of resistance to H_2_O_2_. To test this hypothesis, we selected a serial of H_2_O_2_-resistant mutant strains using a clinically important stressor H_2_O_2_ and identified the highly H_2_O_2_-resistant mycobacterial strain mc^2^51 (Li et al., 2014b). Compared to the wild-type *M. smegmatis* mc^2^155 strain, the minimum inhibitory concentration (MIC) of H_2_O_2_ was more than 80-fold higher, which the MIC of mc^2^51 to H_2_O_2_ was 3.125 mM, similar to that of *M. bovis* BCG (0.625 mM) (Li et al., 2014a) and *M. tuberculosis* (0.625 mM), while that of mc^2^155 was 0.039 mM (Li et al., 2014b). The mc^2^51 exhibited a slow growth rate similar to *M. tuberculosis*.

In this study, we first showed that H_2_O_2_-resistant mc^2^51 had a growth advantage both in mice and macrophages compared with wild-type mc^2^155, which indicated that the higher resistance to H_2_O_2_ of mycobacteria is related to higher virulence. Similar to *M. tuberculosis* that can survival under hypoxic in the lungs and resuscitate under appropriate conditions, mc^2^51 presented a growth advantage to recover from dormancy using the Wayne dormancy model. Furthermore, we showed that Fur mutant carrying the A28V point mutation dysregulated *katG* levels, which was the main cause for resistance to H_2_O_2_ and low levels of ATP. Additionally, the redox-related protein, Rv1996, responsible for regulating gene expression, exhibited different phenotypes associated with isoniazid susceptibility in *M. smegmatis* mc^2^155, mc^2^51, *M. bovis* BCG and *M. tuberculosis* mc^2^7000. Thus, the same protein presents different phenotypes under different physiological conditions. Our results suggest that the difference in the corresponding redox status causes the difference in pathogenicity between *M. tuberculosis* and *M. smegmatis*.

## RESULTS

### The *Mycobacterium tuberculosis-like M. smegmatis* mutant strain mc^2^51 displayed improved virulence

Previous studies showed that *M. smegmatis* with the gain of H_2_O_2_-resistance, named mc^2^51, improves growth fitness in mycobacteria under stress (Table 1) (Li et al., 2014a; Li et al., 2014b). Prior to selecting H_2_O_2_-adapted mycobacterial mutants, we first measured the MIC of wild-type mc^2^155 to H_2_O_2_, which was 0.039 mM. We used 0.0293 mM H_2_O_2_ for the initial screening, which was lower than the MIC of mc^2^155 to H_2_O_2_ (Table 1). The whole process was performed as follows: cultures were started from glycerol-frozen stocks and grown to log phase (OD_600_ of 0.6-0.8). Then the culture was diluted 1000 times and grown under 0.0293 mM H_2_O_2_ until the OD_600_ reached logarithmic (log) phase. Cultures were then further diluted 1:1000 and an additional 0.0293 mM of H_2_O_2_ was added to the culture. This process was repeated until the H_2_O_2_ concentrations reached 0.4395 mM. In further rounds of culture, H_2_O_2_ was added in steps of 0.0879 mM, instead of 0.0293 mM until and concentration of 1.5 mM was reached (Supplemental Figure 1 and Supplement Table 1). The actual MIC of H_2_O_2_ of the mutant strain, named mc^2^51, selected at 1.5 mM was 3.125 mM.

**Table. 1.** List of bacteria in this study.

| Strains | Genotype or relevant characteristics | Source or reference | H<sub>2</sub>O<sub>2</sub> (mM) | INH (μg/mL) |
| --- | --- | --- | --- | --- |
| mc<sup>2</sup>155 | *Mycobacterium smegmatis* | Snapper et al., 1990 | 0.039 | 10 |
| mc<sup>2</sup>51 | highly H<sub>2</sub>O<sub>2</sub> resistance, *Mycobacterium tuberculosis-like Mycobacterium smegmatis* | Li et al., 2014a; Li et al., 2014b | 3.125 | 0.1 |
| $\Delta katG$ | *Mycobacterium smegmatis* $\Delta katG::hyg^R$ | This study | N/A | N/A |
| pMV261-*katG*/mc<sup>2</sup>155 | *Mycobacterium smegmatis* harboring pMV261- *katG* | This study | N/A | N/A |
| *mFur*/mc<sup>2</sup>155 | *Mycobacterium smegmatis* *mFur* at T28A site | This study | 0.64 | N/A |
| pMV261/mc<sup>2</sup>155 | *Mycobacterium smegmatis* harboring pMV261 Kan<sup>R</sup> | This study | N/A | 2.5 |
| pMV261-*rv1996*/mc<sup>2</sup>155 | *Mycobacterium smegmatis* harboring pMV261-*rv1996*, Kan<sup>R</sup> | This study | N/A | 10 |
| pMV261/mc<sup>2</sup>51 | highly H<sub>2</sub>O<sub>2</sub> resistance *Mycobacterium smegmatis* harboring pMV261 Kan<sup>R</sup> | This study | N/A | 0.1 |
| pMV261-*rv1996*/mc<sup>2</sup>51 | highly H<sub>2</sub>O<sub>2</sub> resistance *Mycobacterium smegmatis* harboring pMV261-*rv1996*, Kan<sup>R</sup> | This study | N/A | 0.1 |
| pMV261/BCG | *Mycobacterium bovis* BCG Pasteur harboring pMV261, Kan<sup>R</sup> | This study | N/A | 0.05 |
| pMV261-*rv1996*/BCG | *Mycobacterium bovis* BCG Pasteur harboring pMV261-*rv1996*, Kan<sup>R</sup> | This study | N/A | 0.025 |
| pMV261/mc<sup>2</sup>7000 | *Mycobacterium tuberculosis* Δ*panCD* Harboring pMV261, Kan<sup>R</sup> | This study | N/A | 0.05 |
| pMV261-*rv1996*/mc<sup>2</sup>7000 | *Mycobacterium tuberculosis* Δ*panCD* Harboring pMV261-*rv1996*, Kan<sup>R</sup> | This study | N/A | 0.05 |

N/A: Not available

In a previous study, the mutated mycobacterial strain, mc^2^51, evolved into an *M. tuberculosis-like* strain with presenting slow growth and improves growth fitness under stress (Li et al., 2014b), which H_2_O_2_ resistance in mycobacteria linked to virulence. Previous studies including ours have associated isoniazid (INH) with H_2_O_2_-resistance (Hu et al., 2021; Timmins and Deretic, 2006). The MIC of isoniazid (INH) for mc^2^51 was dramatically reduced to ~ 1% of the MIC for mc^2^155 (Table 1). To test our hypothesis that the H_2_O_2_-resistant strain mc^2^51 would be more persist in the host than its parental strain mc^2^155, we performed non-invasive intranasal infections, which can induce respiratory mucosal immune responses and is a promising way of vaccination for respiratory infections diseases. As shown in Figure 1A, intranasally infection of C57BL/6 mice with ~1×10^7^/50 μL and bacterial colony-forming units (CFUs) were counted 1 day after infection. Compared to mc^2^155 with the percentage of survival in the infected lung of 0.03653 ± 0.01462 % (n = 3), the mc^2^51 strain with that of 0.7153 ± 0.2597 % (n = 3) had a significantly higher survival percentage (p = 0.0451) (Figure 1B). After infection with live mycobacteria, the bacilli entries into alveolar macrophages (AMΦs) and persist within them. To exclude the possibility that the higher CFUs were caused by the larger number of infected bacilli being captured by AMΦs, macrophage-killing assays were explored using the THP-1 cell line. The survival percentage of mutant mc^2^51 is 0.3158 ± 0.1502 (n = 9) while survival of wild type mc^2^155 was 0.01921 ± 0.01223 (n = 6) (p = 0.0003) (Figure 1C). These results indicated that mc^2^51 exhibited enhanced virulence.

**Figure 1.**
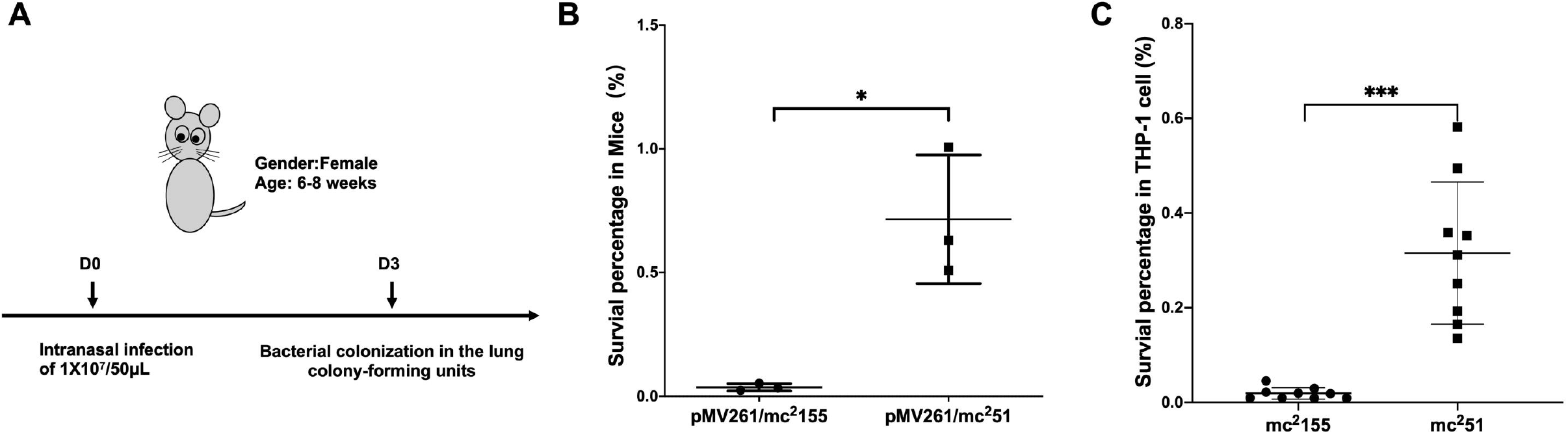
*M. smegmatis* mc^2^51 displayed improved virulence in mice and THP-1 cells. **(A)** The schematic diagram of the mice infection. C57BL/6 mice were intranasally infected as described in Materials and Methods with ~1×10^7^/50 μL of *M. smegmatis* mc^2^155 or mc^2^51. At 3rd day postinfection, the whole lung homogenates were plated to determine bacterial numbers. **(B)** The H_2_O_2_-resistant mutant strain mc^2^51 has a significantly higher bacterial load in the infected lung. The lung burdens after infection with mc^2^155 (black circles) or mc^2^51(black squares) after 3-day postinfection were measured and the survival percentage was calculated as described in Materials and Methods. Each group consisted of 3 mice. * *p* < 0.05. **(C)** The H_2_O_2_-resistant mutant strain mc^2^51 has a survival advantage over wild-type mc^2^155 in the macrophage like cell line THP1.THP1 cells were infected with mc^2^155 (black circles) or mc^2^51(black squares) at a multiplicity of infection of 1 h after infection, THP cell lysates were collected and the intracellular bacilli were measured and the survival percentage was calculated as described in Materials and Methods. Each group consisted of 9 repeats. \*\*\**p* < 0.001.

### Mutant *fur* altered the intracellular redox state and was conductive to latency and resuscitation via modulation of *KatG* levels

*M. tuberculosis* has evolved to survive in hypoxic conditions. Over more than 100 years of research, *M. tuberculosis* has been confirmed to be an obligate aerobic bacterium that cannot replicate under hypoxic conditions. However, *M. tuberculosis* has incredible survivability in long-term anaerobic environments. Evidence suggests that *M. tuberculosis* has the ability to reduce the respiratory system to low levels and maintain vitality (Loebel et al., 1933). Due to respiratory depression, ATP is maintained at low levels, which guarantees minimal metabolic activity to ensure membrane integrity under hypoxic conditions. We first compared the survival of mc^2^51 and mc^2^155 strains under hypoxic conditions. We established a Wayne dormancy model (Wayne and Hayes, 1996), consisting of mycobacterial strain cultures grown in 7H9 medium to an OD_600_ of 1.0, which were then transferred to anaerobic tubes containing 1×10^6^ cells/mL with a headspace ratio of the culture system of 0.5. Methylene blue (1.5 mg/L) was added as an indicator of oxygen status. The OD_600_ and CFUs at different indicated times and the color transition time were measured. The indicator of mc^2^51 has become colorless on the 6th day (141 hours), on the contrary, the indicator of mc^2^155 was still in blue at this time (Figure 2A). For mc^2^155, the indicator became colorless in about 8 days (189 hours), indicating that it has entered an anaerobic state. The mc^2^51 entered the anaerobic state faster in the later stages, and the number of viable bacteria in the anaerobic state was significantly higher than mc^2^155 (Figure 2B and 2C).

**Figure 2.**
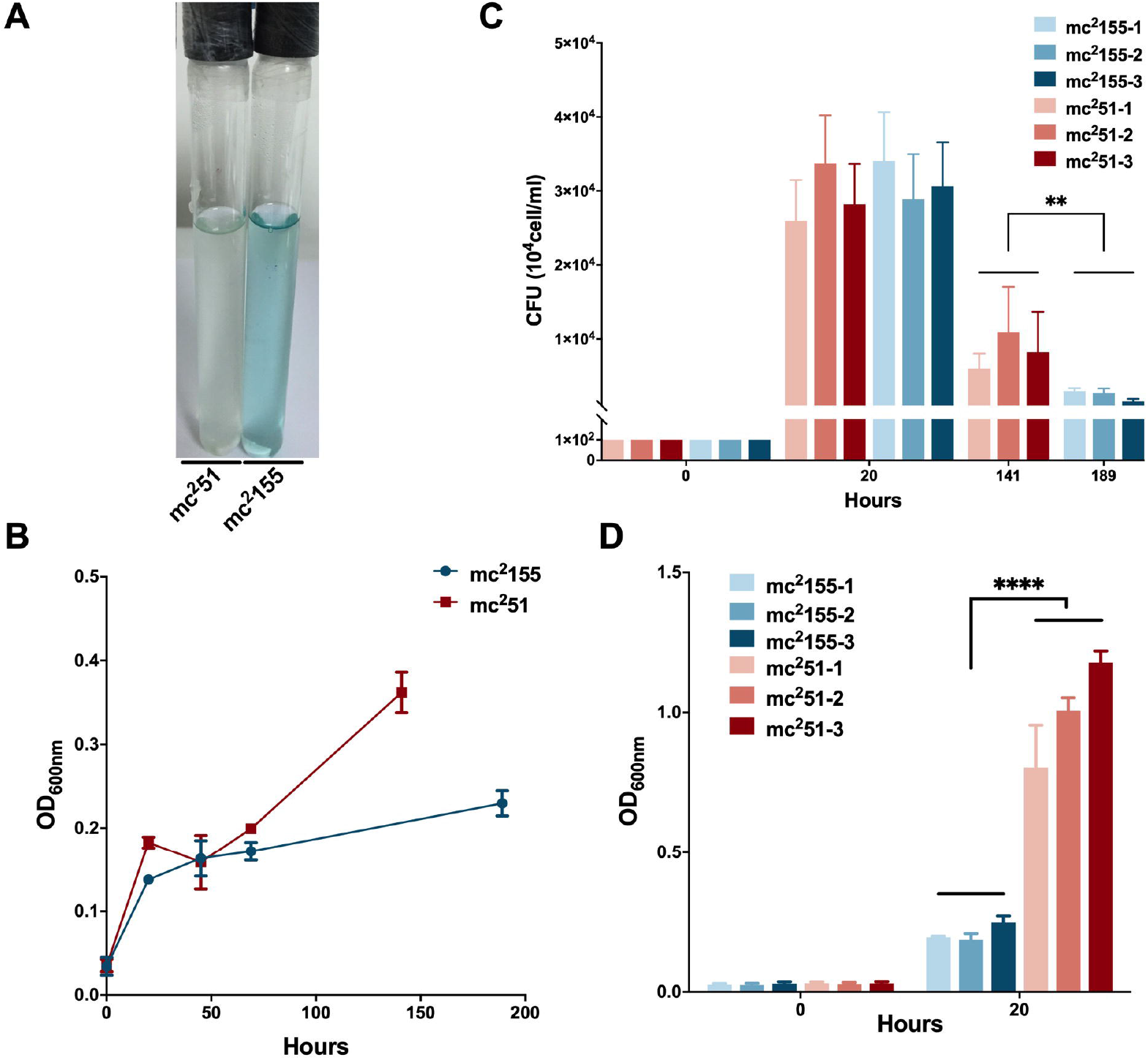
Survival in hypoxic environments and resuscitation of *M. smegmatis* mc^2^155 and mc^2^51. **(A)** The oxygen tension indicator Methylene blue of mc^2^51 culture (left) changes to colorless on day 6th, on the contrast, the indicator of mc^2^155 culture (right) maintains blue in the Wayne dormancy model. The cultures were initially inoculated in anaerobic tubes at OD_600_ of 0.01 with 0.5 head-space ratio. The cultures were stirred at 120 rpm. Methylene blue (1.5 mg/L) was used as an oxygen tension indicator. Methylene blue changes in color from blue to colorless under reducing conditions. Data present results of three biological replicates. Growth rates of strains mc^2^155 (navy blue) and mc^2^51 (brown) were measured by measuring the OD_600_ (B) and by determination of CFUs (C) after plating on 7H10. Data are presented as the mean ± standard deviation of three independent replicates. \*\**p* < 0.01. **(D)** The H_2_O_2_-resistant mutant strain mc^2^51 shows a growth advantage of recovering from dormancy in an anaerobic state over mc^2^155. After the indicator methylelen blue in the culture of Wayne dormancy model became coloress, cultures of mc^2^155 (navy blue) or mc^2^51 (brown) were collected and reinoculated into 7H9 combinated wih heart infusion medium at OD_600_ of 0.01 and aerobic shaked at 200 rpm. The CFUs were determined after 20 h post-inoculation. Data are presented as the mean ± standard deviation of three independent replicates. **** *p* < 0.0001

For the activation experiment, bacteria cultured in the anaerobic conditions were collected and diluted to 1×10^6^ cells/mL in 7H9 and the brain-heart infusion medium and were then grown under aerobic conditions. The three independent mc^2^51 clones were set and each clone set up 3 replicates, all of which were grown better than the 3 independent clones of mc^2^155 (Figure 2D). Thus, mc^2^51 had the growth advantage of recovering from the dormancy of the anaerobic state over mc^2^155. We also measured the intracellular ATP levels of mc^2^51 and mc^2^155. As expected, the abundance of ATP in mc^2^51(0.1885 ± 0.0481 μM/mg) was lower than that of mc^2^155 (0.8138 ± 0.1324 μM/mg) (Figure 3B).

**Figure 3.**
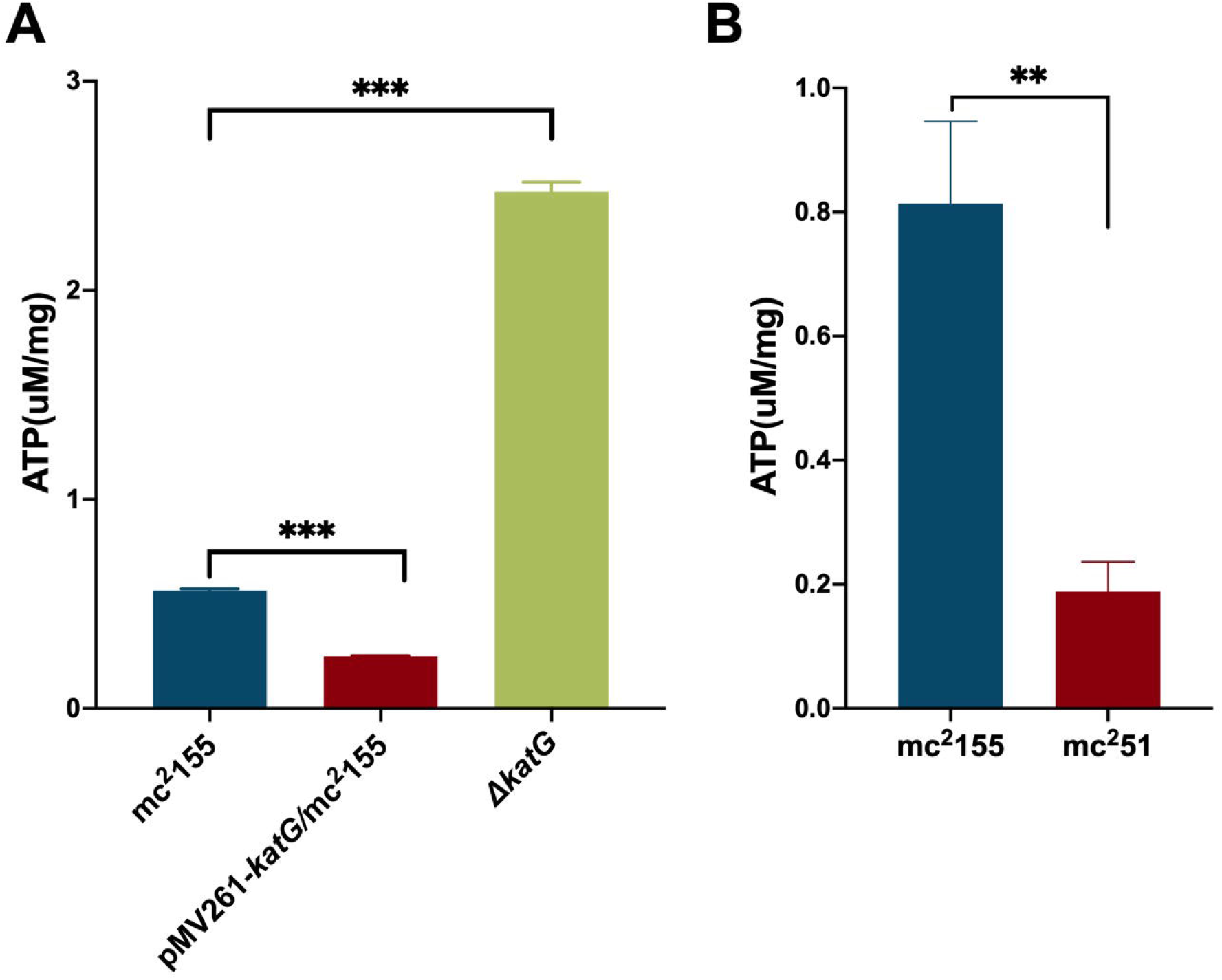
The ATP content in mc^2^51 is similar with pMV261-*katG*/mc^2^155. **(A)** The detection of ATP content in mc^2^155, mc^2^155Δ*katG* and pMV261-*katG*/mc^2^155. The cultures of mc^2^155 (navy blue), pMV261-*katG*/mc^2^155 (brown) and *ΔkatG* (olive drab) at OD_600_ of 0.8 were collected. ATP levels were measured in relative light unit (RLU) using Cytation 3 Cell Imaging Multi-Mode Reader. The corresponding ATP concentrations were calculated according to the ATP standard curve and further converted to μM/mg protein. Data are presented as the mean ± standard deviation of three independent replicates. *** *p* < 0.001 **(B)** The intracellular ATP content of mc^2^51 and mc^2^155. The cultures of mc^2^155 (navy blue) or mc^2^51(brown) at OD_600_ of 0.8 were collected. ATP levels were measured in relative light unit (RLU) using Cytation 3 Cell Imaging Multi-Mode Reader. The corresponding ATP concentrations were calculated according to the ATP standard curve and further converted to μM/mg protein. Data are presented as the mean ± standard deviation of three independent replicates. \*\**p* < 0.01

We previously performed whole-genome sequencing to compare differences in mc^2^155 and mc^2^51 at the genome level. Whole-genome sequencing revealed that there were 29 single nucleotide polymorphisms (SNPs) in mc^2^51. All 29 SNPs were cloned and each was transformed into mc^2^51 strains, only the *fur* (*msmeg_3460*), located upstream of *msmeg_3461*, encoding KatG (Figure 4A), could restore the resistance phenotype of H_2_O_2_ (Li et al., 2014b). The *fur* encoded protein Fur is negatively regulated *katG* expression (Pym et al., 2001). The A28V Fur mutation (mFur) in mc^2^51 may also affect the expression of *katG*. To verify this hypothesis, we examined the binding of mFur to the target DNA (the promoter region of the *fur*) using electrophoretic mobility shift assays (EMSA). Compared to wild-type Fur protein, EMSA showed that mFur decreased DNA binding (Figure 4B), which resulted in the *katG* transcription dysregulation by mFur. In addition, the RNA of mc^2^155 and mc^2^51 was extracted and quantified. The results showed that compared to the mc^2^155 strain, the expression of the catalase-peroxidase KatG encoding gene *katG* of the mc^2^51 strain was significantly up-regulated to ~ 61.82-fold that of the wild-type strain. Taken together, mFur increases KatG protein level in mc^2^51. KatG is a dual-enzyme for catalase and peroxidase, which hydrolyzes ROS (Ng et al., 2004). Thus, the mc^2^51 may maintain ATP at lower levels through KatG, compared to mc^2^155 levels; that is, the abundance of KatG may affect the mycobacterial redox state and thus change the susceptibility to H_2_O_2_. We then constructed the Δ*kalG* (mc^2^155 with knockout *katG*), pMV261-*katG*/mc^2^155 (mc^2^155 with overexpression of *katG*) strains, and their respective ATP content was tested. As shown in Figure 3A, the KatG level negatively correlated with the ATP level. Furthermore, we constructed the specific site mutant of the *fur* gene (*mfur*) in wild-type mc^2^155 causing an amino acid change of A28V of Fur (Figure 5A and 5B) by using recombination protein gp61 from Che9c mycobacteriophage (van Kessel et al., 2008), to construct mc^2^155-*mfur* (Table 1). As we expected, the Fur mutation at A28V induced high resistance to H_2_O_2_ with the MIC of H_2_O_2_ in mc^2^155-*mfur* is 0.64 mM. We showed that the point mutation of *fur* dysregulation of *katG* expression, which is a major factor leading to the phenotype H_2_O_2_-resistance.

**Figure 4.**
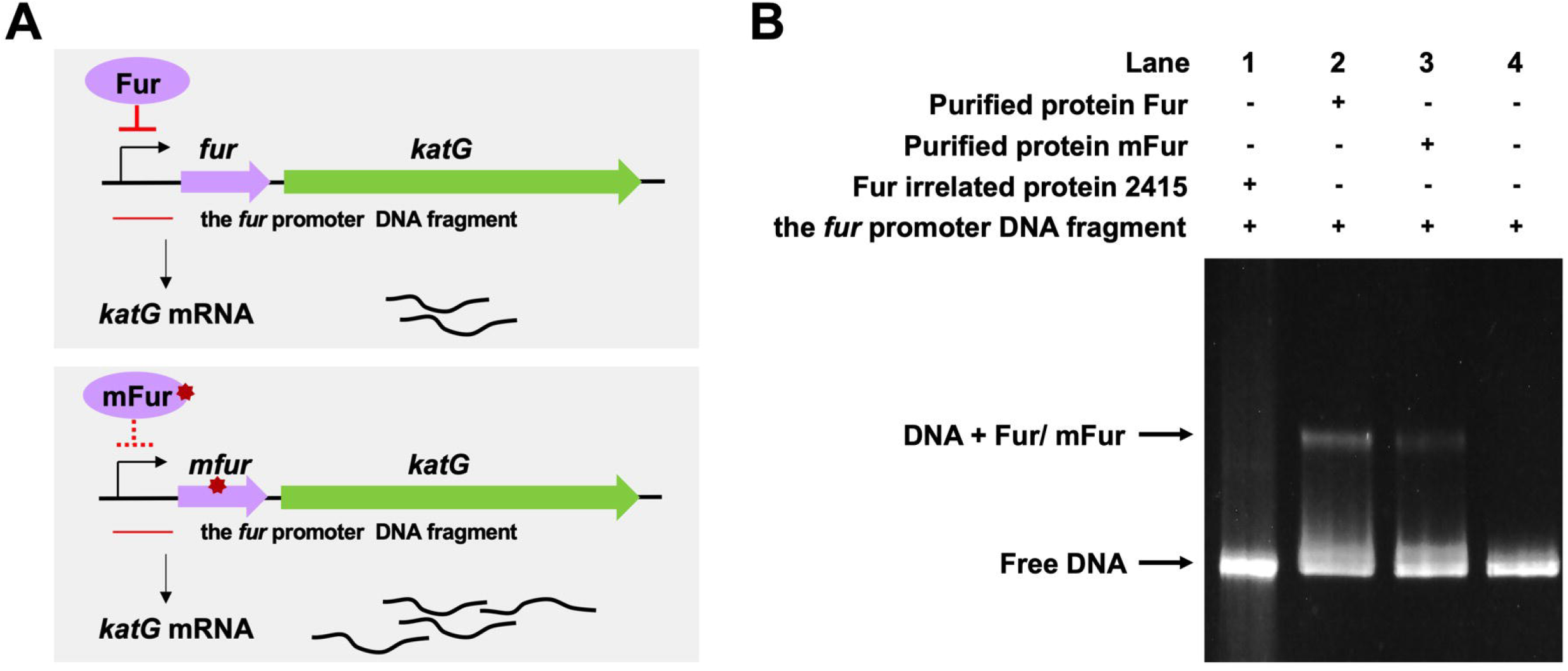
The A28V Fur mutant protein decreased DNA binding to the *fur* promoter. **(A)** Genetic organization of the *fur-katG* and the schematic diagram of Fur negatively regulation of *katG* (Upper panel). Genetic organization of the *mfur-katG* and the schematic diagram of mFur resulted in derepression of *katG* with increasing *katG* mRNA (Bottom panel). The red line indicates the *fur* promoter DNA fragment. **(B)** Electrophoretic mobility shift assays (EMSA) of the binding of Fur/mFur protein to *fur* promoter DNA fragment. Purified MSMEG_2415 protein (2415), irrelated to Fur, expressed in *E. coli* was run in the first lane of an 4-20% polyacrylamide gel. Gel shift caused by Fur (lane 2) and mFur (lane 3). The image shown is representative of at least 3 experiments.

**Figure 5.**
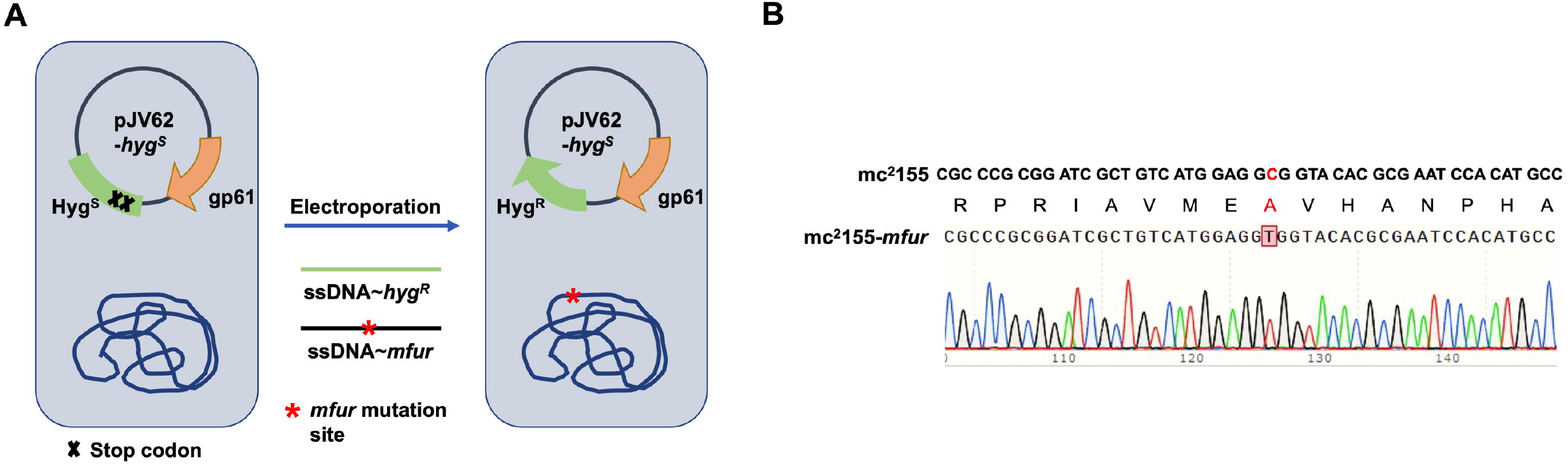
Generation of a point mutation furA28V in fur on chromosome in *M. smegmatis*. **(A)** Strategies for recombineering. The site-directed mutagenesis of fur was obtained using Phage Che9c gp61-mediated recombination. Plasmids pJV62-*hyg^s^* express Che9c gene product gp61, to facilitate single-stranded DNA carrying the point mutation recombination. The clones carrying the mutated site were selected on 7H9 with HygR and KanR. (B) The sequencing result of a point mutation in fur. Sanger sequencing showed that the nuclear acid point mutation site c/t, resulting in the protein switch from A to V at site 28th.

### The same protein performs different functions in different redox states

As a successful human pathogen, *M. tuberculosis* has unique respiration properties. *M. tuberculosis* excretes alkaline supernatants, which is in contrast, to other strains that excrete acidic supernatants (Merrill, 1930). The difference between secreted compounds with different acid-base properties suggested that *M. tuberculosis* has a distinctive redox state. As shown in Table 2, the comparative genomic analysis shows that PhoPR, DosR/S/T, identified as virulence factors of *M. tuberculosis*, are present in *M. smegmatis*. The signaling transduction systems such as the two-component systems and the sigma factors of *M. tuberculosis* are homologous in *M. smegmatis* (Table 3). Different phenotypes might be due to different redox states. Thus, we considered that the same redox-regulated related protein might perform different functions in different redox states. To test this hypothesis, we examined the biological function of a universal stress protein Rv1996 that increases the expression of *KatG*, in various mycobacterial strains (Hu et al., 2015). Both previous studies and our studies have linked isoniazid action with redox states and H_2_O_2_ resistance is negatively correlated to INH susceptibility in mycobacteria (Bhaskar et al., 2014; Hu et al., 2015; Vilcheze et al., 2017). We used INH as a chemical probe for monitoring mycobacterial redox states and measured the MICs of INH to corresponding mycobacterial strains (Figure 6). As predicted, the MICs of INH differed across the tested mycobacterial strains: the MIC of INH in pMV261-*rv1996*/mc^2^7000 was equal to that of pMV261/mc^2^7000; the MIC of INH in pMV261-*rv1996*/BCG was lower than that in pMV261/BCG; the MIC of INH in pMV261-*rv1996*/mc^2^51 was equal to that of pMV261/mc^2^51; and in mc^2^155, the opposite results were observed with the MIC of INH in pMV261-*rv1996*/mc^2^155 being lower than that of INH in pMV261/mc^2^155 (Figure 6).

**Figure 6.**
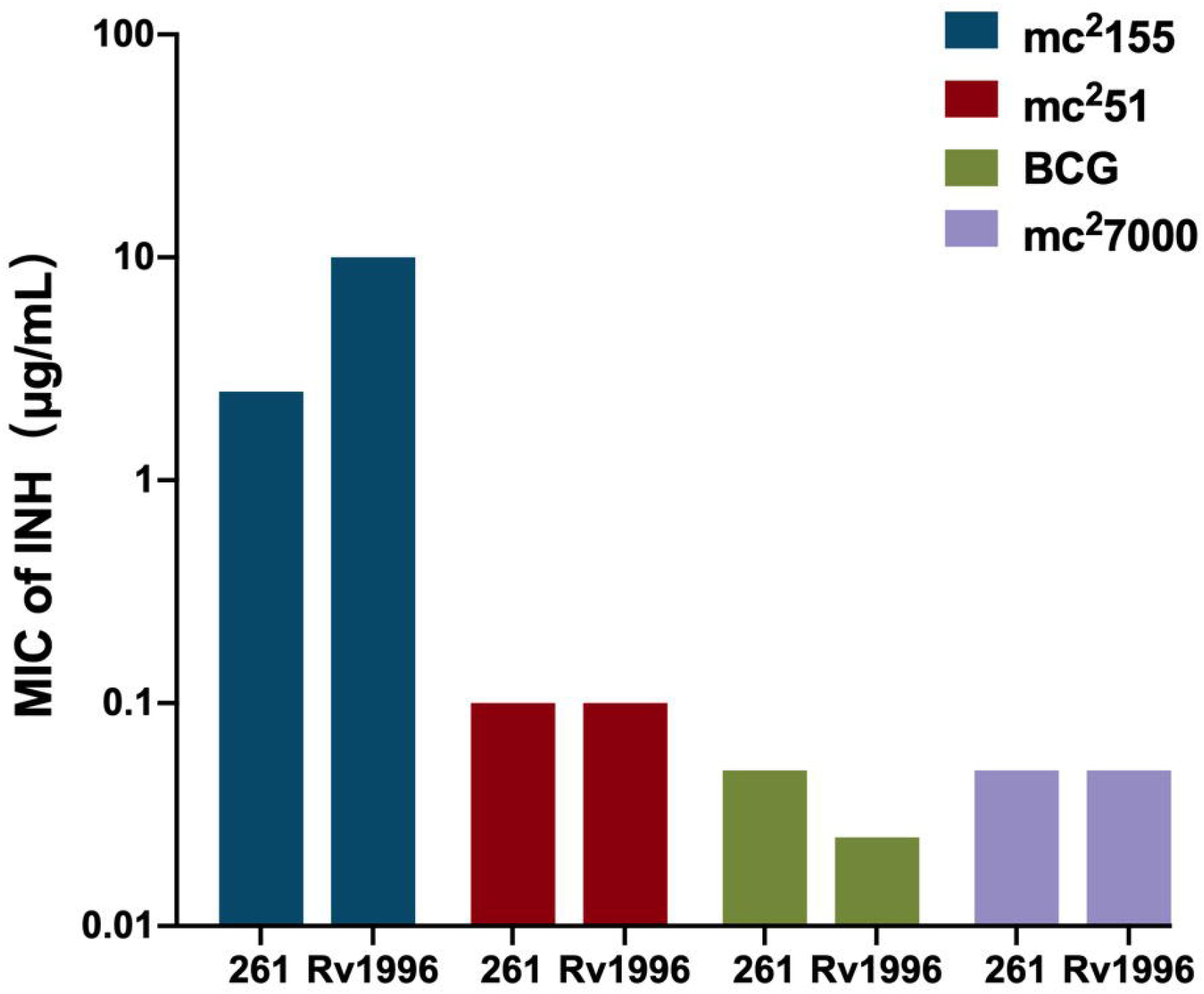
Comparison of isoniazid susceptibility in different mycobacterial strains harboring *pMV261-rv1996* and the corresponding control strain harboring pMV261. The minimum inhibitory concentrations (MICs) of isoniazid were determined by inoculating each bacterial strain in 7H9 containing serially isoniazid (INH). The values of MIC were recorded. 261 present pMV261. Rv1996 present pMV261-*rv1996*. The image represents the results of three independently repeat.

**Table.2.** Conservation of *M. tuberculosis* H37Rv TCSS in *M. smegmatis*.

| Gene | *M. tuberculosis* Rv# | Mtb | Msm | Reference |
| --- | --- | --- | --- | --- |
| RegX3-SenX3 | Rv0491- Rv0490 | + | + | James et al., 2012; Parish et al., 2003b; Rifat and Karakousis, 2014 |
| HK1-HK2- tcrA | Rv0600c-Rv0601c-Rv0602c | * | – | Shrivastava and Das, 2007 |
| PhoP-PhoR | Rv0757- Rv0758 | + | + | Walters et al., 2006 |
| NarL-NarS | Rv0844c-Rv0845c | + | + | Schnell et al., 2008 |
| PrrA-PrrB | Rv0903c- Rv0902c | + | + | Arora et al., 2021; Nowak et al., 2006 |
| MprA-MprB | Rv0981- Rv0982 | + | + | He and Zahrt, 2005; Sureka et al., 2007 |
| KdpD-KdpE | Rv1028c-Rv1027c | + | + | Parish et al., 2003a; Steyn et al., 2003 |
| TrcR-TrcS | Rv1032c- Rv1034c | + | + | Haydel et al., 2002 |
| MtrA-MtrB | Rv3245c-Rv3247c | + | + | Fol et al., 2006; Li et al., 2010; Plocinska et al., 2012 |
| TcrX-TcrY | Rv3765c-Rv3764c | + | + | Bhattacharya et al., 2010 |
| PdtaS-PdtaR | Rv3220c-Rv1626 | + | + | Boshoff et al., 2004; Shrivastava and Das, 2007 |

Msm, *Mycobacterium smegmatis*; Mtb, *Mycobacterium tuberculosis* CDC1551;
+, Genes encoding the sensor kinase and the response regulator are present and genetically linked;
–, genes encoding both the sensor kinase and the response regulator are absent;
\*, Genes encoding the two sensor kinases have been fused, and the gene encoding this fused sensor
kinase is genetically linked to the response regulator.

**Table. 3.** Sigma factor genes in mycobacteria.

| Sigma | *M. tuberculosis*<br>Rv# | Mtb | Msm | Reference |
| --- | --- | --- | --- | --- |
| SigA($\sigma^A$) | Rv2703 | + | + | Gomez et al., 1998 |
| SigB($\sigma^B$) | Rv2710 | + | + | Lee et al., 2008b; Mukherjee and Chatterji, 2005 |
| SigC($\sigma^C$) | Rv2069 | + | – | Sun et al., 2004 |
| SigD($\sigma^D$) | Rv3414c | + | + | Calamita et al., 2005; Raman et al., 2004 |
| SigE($\sigma^E$) | Rv1221 | + | + | Song et al., 2008 |
| SigF($\sigma^F$) | Rv3286c | + | + | Rodrigue et al., 2007 |
| SigG($\sigma^G$) | Rv0182c | + | + | Lee et al., 2008a |
| SigH($\sigma^H$) | Rv3223c | + | + | Song et al., 2008 |
| SigI($\sigma^I$) | Rv1189 | + | – | |
| SigJ($\sigma^J$) | Rv3328c | + | + | Homerova et al., 2008 |
| SigK($\sigma^K$) | Rv0445c | + | – | Veyrier et al., 2008 |
| SigL($\sigma^L$) | Rv0735 | + | + | Hahn et al., 2005 |
| SigM($\sigma^M$) | Rv3911 | + | + | Agarwal et al., 2007; Raman et al., 2006 |

+, presence of the gene;
–, absence of the gene.

## DISCUSSION

Understanding how the *M. tuberculosis* evolved into a professional pathogen is of benefit to the study of its pathogenesis and design of vaccines. The combination of experimental evolution and whole-genome sequencing provides a powerful method for identifying adaptive mutations and elucidating the specific genotype–phenotype relationship (Elena and Lenski, 2003; Lenski, 2017). Historically, the most successful example of continuous selective cultures is *M. bovis BCG*, the only anti-TB vaccine, which was attenuated after 13 years of continuous *in vitro* passages of *M. bovis BCG*. We previously used a similar adaptive evolution strategy to select H_2_O_2_-resistant *M. smegmatis* strains by using a clinically key stressor H_2_O_2_ (Li et al., 2014b). Preliminary results showed that the mc^2^51 strain was highly resistant to H_2_O_2_ and had greater susceptibility to INH, compared to mc^2^155. The mc^2^51 phenotype showed an *M. tuberculosis*-like *M. smegmatis* phenotype. Altogether, the mutant *M. smegmatis* mc^2^51 exhibited higher virulence.

The whole-genome sequencing showed the presence of gene mutations in *fur*, and the mutant Fur resulted in *katG* levels (Figure 4B). In the Wayne dormancy model, mc^2^51 shows a growth advantage of recovering from dormancy under anaerobic conditions over mc^2^155 (Figure 2B). In parallel, a high level of *katG* in mc^2^51 is accompanied by lower ATP levels, which implied mc^2^51 exhibited at a lower level of respiration (Figure 3B). Moreover, we showed that a redox-related protein Rv1996 exhibits a different phenotype under different specific redox states in *M. smegmatis* mc^2^155, mc^2^51, *M. bovis* BCG and *M. tuberculosis* mc^2^7000 (Figure 6). This study indicated that the same genotype presents different phenotypes under different physiological conditions. We at least partially explain why *M. smegmatis* and *M. tuberculosis* have similar virulent factors, including two-component system, and sigma factors (Table 2 and 3), but *M. smegmatis* is a non-pathogen and *M. tuberculosis* is a pathogen.

*M. tuberculosis* is a successful human pathogen. It is considered to be derived from the environment (Gutierrez et al., 2005; Wolfe et al., 2007) and has adapted to the immune environment of the human body through long-term evolution. Its successfully established infection is partially attributed to its survival capacity and persistence in macrophages (Podinovskaia et al., 2013). To defend against mycobacterial infection, the host produces ROS, as an important innate defense mechanism. Consequently, *M. tuberculosis* has evolved a hierarchy and unique antioxidant function and maintains a low level of respiration, manifested by slow growth and persistence in the host. In contrast, *M. smegmatis* is present in the soil, which is a totally different environment from the host (Zhang and Furman, 2021). In Table 2, we show that *M. tuberculosis* and *M. smegmatis* have similar genotypes; however, they show different phenotypes, in terms of INH susceptibility, H_2_O_2_ resistance, and virulence. We believe that this striking difference is due to H_2_O_2_ resistance. The selected resistance to H_2_O_2_ of mc^2^51 shows improved virulence in both the macrophage-killing assay and in an animal model (Figure 1). In fact, several studies have shown that abiotic stress can improve the virulence phenotype of bacterial pathogens (Li et al., 2021; Sundberg et al., 2014). Our study also supports of sit-and-wait hypothesis (Wang et al., 2017), that is, bacterial environmental abiotic stress and virulence evolution. In addition, this study also suggests that we can use mc^2^51 as a model strain, replacing mc^2^155, to study the regulation of redox homeostasis of *M. tuberculosis*.

We previously sequenced the whole genome of mc^2^51 strain (Li et al., 2014a), and identified 29 SNPs, compared to mc^2^51. Confirmed with our previous study (Li et al., 2014b), we found that only the *fur* gene can partially complement the resistant phenotype. This suggested that the *fur* mutation is a major mutation site that causes mc^2^51 to be highly resistant to H_2_O_2_, and other mutation sites may correspond to the growth fitness compensation mutation in mc^2^51. We also mutated the *fur* in wild-type mc^2^155 by genome editing and produced a phenotype similar to mc^2^51, which is highly resistant to H_2_O_2_ (Table 1). Large-scale whole-genome sequencing studies on the evolutionary history of tuberculosis also show that key tract mutations at the transcription site will have a critical impact on the particular phenotype (Gagneux, 2018). For example, the change of PhoP in BCG allows infection with bovine pathogenic bacteria capable of infecting humans (Broset et al., 2015; Gonzalo-Asensio et al., 2014). This study reminds us that when designing vaccines, greater attention should be paid to regulators, which may be more efficient targets. *M. smegmatis* is an effective vaccine for TB and HIV (Kim et al., 2017; Sweeney et al., 2011). The disadvantage of *M. smegmatis* as a vaccine vector is that its transient infection and difficult to establish a persistent infection and produce adaptive immunity. The *M. tuberculosis*-like mutant *M. smegmatis* mc^2^51 may be developed as a vaccine vector.

By comparing the survival of mc^2^155 and mc^2^51 in the Wayne dormancy animal model, we also found that low respiratory levels are beneficial for survival under anaerobic conditions and resurrection (Figure 2). In the future, we plan to use these strains to compare physiological indicators such as NADH/NAD+, NADPH/NADP+, and ATP, to further understand the mechanisms underlying *M. tuberculosis* resurrection. This study provides insight into H_2_O_2_ resistant mechanisms in mycobacteria and has important implications for linking mycobacteria redox capacity and persistence infection in mice.

## MATERIALS AND METHODS

### Strains and growth conditions

The H_2_O_2_-resistant *Mycobacterium smegmatis* strain mc^2^51 was screened in the Mi lab (Li et al., 2014b). *Mycobacterium tuberculosis* Δ*panCD* (named mc^2^7000) (Sambandamurthy et al., 2002) was kindly gifted from J Deng. The *M. smegmatis* wild-type mc^2^155, mutant strain mc^2^51, *M. bovis*

BCG Pasteur *and M. tuberculosis* mc^2^7000 were cultured in Middlebrook 7H9 (Becton Diskinson Sparks, MD, USA) supplemented with ADS (10 % albumin, dextrose and saline), 0.05% Tween 80 (Sigma, St. Louis, MO, USA) and 0.5% glycerol (Beijing Modern Eastern Fine Chemical Co. Ltd., Beijing, China) for liquid culture and Middlebrook 7H10 (Becton Diskinson Sparks, MD, USA) supplemented with ADS for bacterial colonies culture. The colony-forming units (CFUs) of mycobacterial strains were determined by plating serial dilutions of cultures on Middlebrook 7H10 agar plates and incubating at 37°C in an atmosphere of 5% CO_2_ for the indicated time. For mc^2^7000 culture, panthothenate (24 mg/L) was added. When required, kanamycin (25 mg/L, Amresco, USA) and hygromycin (50 mg/L, Sigma, USA) was added. All bacterial strains used in this study were listed in Table 1.

### Determine of MIC to Isoniazid and H_2_O_2_

The susceptibility of isoniazid (INH) or H_2_O_2_ of mycobacteria was evaluated using the modified broth microdilution method (Franzblau et al., 1998). Briefly, INH or H_2_O_2_ was serial diluted using 7H9 medium. The diluted fold was 1.25- or 2-fold, when required. Then 40 μL of diluted INH or H_2_O_2_ were mixed with 40 μL of mycobacterial suspension with 1×10^7^ cells/mL in each well of 96-well microtiter plates, and then incubated at 37°C for the indicated days. As an indicator, 0.02% resazurin was added to individual samples and the color switches from blue to pink were recorded after 4 h. All the experiments were performed in triplicate. The abundance of the cultures was measured using a microplate reader (FLUOstar OPTIMA, BMG Labtech). A difference of 2 serial dilutions or more indicated a significant difference in the INH or H_2_O_2_ susceptibility of bacterial strains.

### Mice infection

Female pathogen-free C57BL/6 mice (aged 6–8 weeks old) were purchased from Vital River (Beijing, China). For the intranasal infection of *M. smegmatis*, mice were anesthetized by intraperitoneal injection of pentobarbital sodium (60 mg/kg), and ~10^7^ CFU/50 μL PBS of mc^2^51 or mc^2^155 were introduced dropwise through the nostril of each mouse. The bacterial burden throughout the infection was monitored by collecting whole lung tissue at the indicated times after the mice were euthanized, and serial dilutions were then plated on 7H10 agar plates. The dose of infection was confirmed on day one post-infection by plating whole lung homogenates from three mice on 7H10 agar. The percentage of survival was calculated as (CFUs after infection/CFUs before infection) × 100%.

### Macrophage-killing assay

Human-derived cell line THP1 (ATCC TIB-202) was cultured in RPMI medium with 10% fetal bovine serum (FBS, GIBCO, USA). THP1 cells were activated with 100 ng/ml phorbo-12-myristate-13-acetate (PMA, Sigma, USA) overnight. Infection was carried out at a multiplicity of infection (MOI) of 10 for 1 hour at 37°C and 5% CO_2_ atmosphere. The infected THP-1 cells were washed with RPMI for 3 times and then chased for 1hr. The cells with intracellular bacilli were then washed, and lysed in sterile cold PBST (PBS with 0.05% Tween 20). Lysates were then vortexed, diluted, and plated on 7H10 agar plates as previously described (Chan et al., 1992), The percentage of survival was calculated as (CFUs after infection/CFUs before infection) × 100%.

### Wayne dormancy model and dormancy exit

Mycobacterial strains were cultured under hypoxic conditions as described by Wayne and Hayes (Wayne and Hayes, 1996). Briefly, cultures were initiated at OD_600_ of ~ 0.01 (1 × 10^6^) and incubated in anaerobic tubes with sealed caps. The headspace ratio of the cultures was 0.5. The cultures were stirred using an 8mm Teflon stir bar (Fisher Scientific, USA) at 200 rpm. Methylene blue (1.5 mg/L) was used as an oxygen tension indicator. It changes in color from blue to colorless under low oxygen tension. The color transition time were recorded. All experiments were prepared in triplicate. Growth was monitored by measuring the OD_600_ and by determination of CFUs after plating on 7H10.

The indicator methylene blue in the culture became colorless, indicating that the bacteria entered anaerobic conditions. Then collecting the bacteria in the anaerobic tube, they were washed with culture medium or PBS three times, resuspended in a culture medium (7H9 and brain heart infusion medium), the concentration was adjusted to the same amount of OD_600_ (OD_600_ of ~ 1.0), and diluted it for the CFU count, 1: 100 or 1:50 into fresh culture medium were shaken at 37°C, to monitor the status of the bacteria, and the OD_600_ was measured. Three independent mycobacterial strain clones were set and each clone set up 3 replicates.

### Measurement of intracellular ATP

The ATP Assay Kit was purchased from Beyotime Biotechnology (Beijing, China). The intracellular ATP assay was performed followed the protocol provided by the manufacturer. Briefly, the sample measurements were prepared as follows: cultures of indicated mycobacteria were obtained to an OD_600_ of 0.8. Bacteria were collected by low-temperature centrifugation at the maximum speed and the pellet was washed with precooled PBS buffer 3 times. A 300 μL volume of ATP detection lysate and 0.5 mL volume of glass beads was added for cell lysis. The lysate obtained was centrifuged at low temperature for 5 minutes and the supernatant was placed on ice for later use. The preparation of the standard solution of gradient concentration ATP was performed as follows: the ATP standard solution was serially diluted into 7 concentrations of 10, 3.333, 1.111, 0.37, 0.1234, 0.04115 and 0.01371 μM. and stored on ice for later use. The preparation of the working solution for the detection of ATP was performed as follows: an appropriate amount of ATP detection reagent was prepared according to the number of samples, and then a 90% final volume of ATP detection reagent diluent was added. The prepared working fluid was placed on ice for further use. The determination of the ATP level was performed as follows: (1) the prepared ATP detection working solution was dispensed into 1.5 mL centrifuge tubes, 100 μL per tube and was allowed to incubate at room temperature for 5 minutes to allow full reaction of the ATP in the centrifuge tube; (2) during the test, 20 μL of each sample (standard or total protein sample) was added to a 1.5 mL centrifuge tube containing 100 μL of ATP detection working solution, and was mixed quickly with a pipette, and incubated for 2 seconds to complete the reaction before using Cytation 3 Cell Imaging Multi-Mode Reader to determine the relative light unit (RLU); (3) a standard curve was constructed to measure to determine the concentration of the sample by converting the RLU into an ATP concentration; (4) In order to eliminate the error caused by the difference in the amount of protein during sample preparation, the BCA protein concentration determination kit produce by Beyotime Biotechnology (Beijing, China) was used to determine the protein concentration in the sample. The ATP concentration of ATP was converted to μM/mg protein.

### Electrophoretic mobility shift assay

The coding regions of *fur* and *mfur* were amplified from mc^2^155 and mc^2^51 genomic DNA and cloned into the *Escherichia coli* expression vector pET23b (+) (Novagen, Madison, WI, USA) inframe fused with a C-terminal His_6_-tag sequence to construct the plasmids pET23b-*fur* and pET23b-*mfur*. The final constructs were transformed into BL21 (DE3) for expression and recombinant Fur/mFur proteins were purified using Ni-NTA agarose (Qiagen, California, USA). The proteins induced by the addition of 1 mM IPTG at 16°C for 12 h. Protein purification was performed as described previously (Li et al., 2014c). The protocols of the recombinant protein purification are available on request. The recombinant protein MSMEG_2415 was purified as described previously (Li et al., 2014c) and used as a negative control for EMSA, which MSMEG_2415 is irrelated to Fur. DNA fragment containing the promoter region of *fur* for gel shift experiments were amplified by PCR with specific primers (forward: 5’-CGTTGGAAAACAACCATTGCAAG-3’, reverse: 5’-CATCCGCAGTTGGGCTTCGAAC-3’). Binding reaction mixtures in 20 μL of binding buffer (20 mM Tris HCl pH 8.0, 1 mM dithiothreitol (DTT), 50 mM KCl, 5 mM MgCl_2_) containing 0.15 pmol of the DNA fragment were incubated with purified Fur/mFur protein (0.5 nmol) for 30 min at 30□. Reaction mixtures were loaded on an 4-20% polyacrylamide gel containing 0.5×TBE. Gels were run at 70 V at 4°C for 3 h. The gel was stained with Good-view and photographed for the image.

### Generation of the *katG* knockout and KatG overexpression strains

The knockout *katG* strain was constructed using Mycobacteriophage-based specialized transduction (Li et al., 2014c). The upstream and downstream sequences of *katG* were amplified from *M. smegmatis* genome DNA. The knockout vector was constructed using phAE159 (Hsu and Jacobs, unpublished data). The mycobacteriophage used for knockout was obtained using MaxPlax packaging extract (Epicentre Biotechnologies, Madision, WI, USA) and a *katG* knockout strain was obtained by phage transduction, named Δ*katG*. The KatG overexpression strain was constructed using pMV261 to yield pMV261-*katG* and the constructed plasmid was electroporated into mc^2^155, yielding pMV261-*katG*/mc^2^155. The detailed information for construction of all the mycobacterial strains and primers for plasmid construct are available on request.

### Generation of *fur* point mutation on chromosome in *M. smegmatis*

The single strand (ss) DNA oligonucleotides used for recombineering were ordered from Genewiz company (Suzhou, China). The site-directed mutagenesis of *fur* was obtained using Phage Che9c gp61-mediated recombination (van Kessel et al., 2008). The detailed information on primers for construction of the *fur* mycobacterial strain (named *mc*^2^155-*mfur*) is available on request. The coding region containing *fur* point mutation in genome (encoding mFur) was amplified and sequenced by Genewiz company (Suzhou, China).

### Generation of the *rv1996* overexpression mycobacterial strains

The *rv1996* gene was amplified and constructed and cloned into pMV261 to yield pMV261-*rv1996*. The constructed pMV261-*rv1996* plasmid was transformed into mycobacterial strains, *M. smegmatis* mc^2^155 and mc^2^51, *M. bovis* BCG Pasteur and *M. tuberculosis* mc^2^7000, the corresponding strains named pMV261-*rv1996*/mc^2^155, pMV261-*rv1996*/mc^2^51, pMV261-*rv1996*/BCG, pMV261-*rv1996*/mc^2^7000. The empty vector pMV261 was transformed into corresponding mycobacterial strains, named pMV261/mc^2^155, pMV261 /mc^2^51, pMV261/BCG, pMV261 /mc^2^7000.

### Statistical analysis

Each experiment was carried out at least twice with three-nine mice or samples per group. The CFUs and OD_600_ were analyzed using unpaired t test (Version 8.0 for Windows GraphPad Software). The ATP content were analyzed using ANOVA tests (Version 8.0 for Windows GraphPad Software). \*\*\*\**p* < 0.0001, \*\*\**p* < 0.001, \*\**p* < 0.01 and \**p* < 0.05.

### Animal ethics

This study was performed in strict accordance with the recommendations of Ethics Committee established in the Guide for the Care and Use of Laboratory Animals of the Institute of Microbiology, Chinese Academy of Sciences (IMCAS). The protocol was approved by the Committee on the Ethics of Animal Experiments of IMCAS. The mice were bred under specific pathogen-free conditions at the IMCAS laboratory animal facility. All animal experiments were conducted under isoflurane anesthesia and all efforts were made to minimize suffering.

## Data Availability Statement

The genome sequence of mc^2^51 has been deposited at DDBJ/EMBL/GenBank under the accession no. JAJD00000000 (https://www.ncbi.nlm.nih.gov/nuccore/JAJD00000000).

## Conflict of Interest

The authors declare no conflict of interest.

## Author Contributions

K.M. conceived and designed the experiments; Z.J. and Z.Z. performed the experiments; K.M. wrote the paper, K.M., Z.J. and Z.Z. revised the paper; All authors have read and agreed to the published version of the manuscript.

## Funding/Acknowledgments

This work was supported by grants from the Ministry of Science and Technology of China (2018YFC1603900, 2017YFA0505901 to K.M.), National Natural Science Foundation of China (31970136,32170181 to K.M., 31900117 to L.F.) and International Joint Research Project of the Institute of Medical Science, University of Tokyo (Extension-2019-K3006 to K.M.). We thank J Deng for providing the mycobacterial strain mc^2^7000. We also thank Tong Yin for her help in preparing the experimental materials.

## Notes

### Competing Interest Statement

The authors have declared no competing interest.

