## Supplemental Table 1 for "Experimental Evolution Reveals Redox State Modulates Mycobacterial Pathogenicity"

**Supplemental Figure 1. Evolutionary selection of H_2_O_2_-resistant mutations in *M. smegmatis***

Cultures were started from glycerol-frozen stocks and grown to log phase (OD_600_ of 0.6 - 0.8). Then the cultures were diluted 1:1000 into 5 ml of 7H9 media containing 10% ADS. Hydrogen peroxide (H_2_O_2_) was then added to a concentration of 0.0293 mM and cultures were grown until the OD_600_ reached log phase. Cultures were then further diluted 1:1000 and an additional 0.0293 mM of H_2_O_2_ was added to the culture. This process was repeated until the H_2_O_2_ concentration reached 0.4395 mM. In further rounds of culture, H_2_O_2_was added in steps of 0.0879 mM, instead of 0.0293 mM until an H_2_O_2_ concentration of 1.5 mM was reached. To ensure that the H_2_O_2_-resistant phenotype was caused by a chromosomal mutation, selected cultures were sub-cultured for 10 generations and then streaked on plates to obtain single colonies. The distinctive single colonies were then inoculated in liquid culture and actual MIC of H_2_O_2_ was determined.

**Supplemental Table 1.** **The selection H_2_O_2_ concentrations and growth time**

| **Days** | **Concentration of H_2_O_2_ (mM)** | **Growth time (Day)** | **Number of generations** |
| --- | --- | --- | --- |
| **2** | **0.029** | **2** | **1** |
| **4** | **0.059** | **2** | **2** |
| **7** | **0.088** | **3** | **3** |
| **11** | **0.117** | **4** | **4** |
| **12** | **0.147** | **1** | **5** |
| **16** | **0.176** | **4** | **6** |
| **19** | **0.205** | **3** | **7** |
| **21** | **0.234** | **2** | **8** |
| **24** | **0.264** | **3** | **9** |
| **28** | **0.293** | **4** | **10** |
| **30** | **0.322** | **2** | **11** |
| **32** | **0.352** | **2** | **12** |
| **33** | **0.440** | **1** | **13** |
| **34** | **0.527** | **1** | **14** |
| **36** | **0.615** | **2** | **15** |
| **38** | **0.703** | **2** | **16** |
| **40** | **0.791** | **2** | **17** |
| **41** | **0.879** | **1** | **18** |
| **43** | **0.967** | **2** | **19** |
| **45** | **1.055** | **2** | **20** |
| **48** | **1.143** | **3** | **21** |
| **50** | **1.231** | **2** | **22** |
| **51** | **1.319** | **1** | **23** |
| **53** | **1.406** | **2** | **24** |
| **54** | **1.494** | **1** | **25** |
