## Supplementary figures and images for "Experimental Evolution Reveals Redox State Modulates Mycobacterial Pathogenicity"

### Supplemental Figure 1

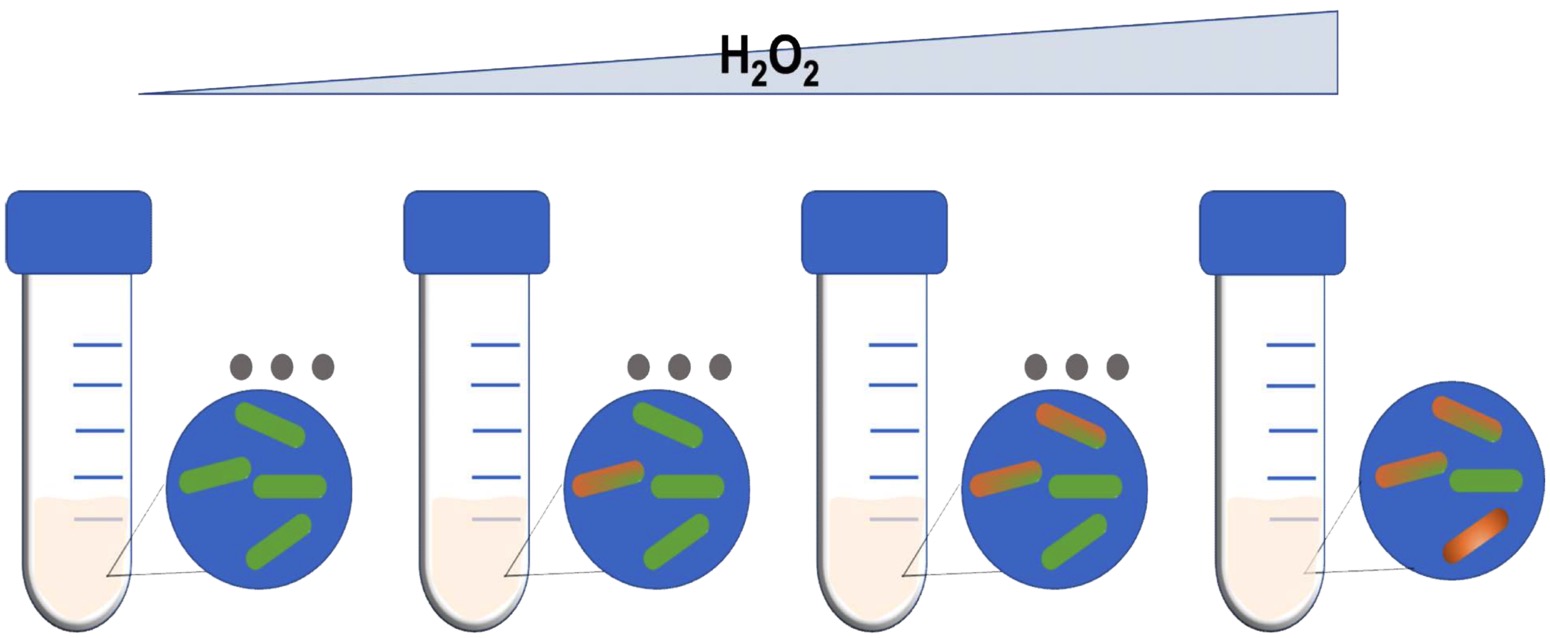
